# A homing rescue gene drive with multiplexed gRNAs reaches high frequency in cage populations but generates functional resistance

**DOI:** 10.1101/2023.11.29.569221

**Authors:** Jingheng Chen, Shibo Hou, Ruobing Feng, Xuejiao Xu, Nan Liang, Jackson Champer

## Abstract

CRISPR homing gene drive is a potent technology with considerable potential for managing populations of medically and agriculturally significant insects. It induces a bias in the inheritance of the drive allele in progeny, rapidly spreading desired genes throughout the population. Homing drives operate by Cas9 cleavage followed by homology-directed repair, copying the drive allele to the wild-type chromosome. However, resistance alleles formed by end-joining repair pose a significant obstacle to the spread of the drive. To address this challenge, we created a homing drive targeting the essential but haplosufficient *hairy* gene. Our strategy involves spreading the drive construct through the homing process, eliminating nonfunctional resistance, which are recessive lethal, while rescuing drive-carrying individuals with a recoded version of *hairy*. This strategy eliminates resistance more slowly than a previous strategy targeting haplolethal genes, but it may be easier to construct in non-model organisms. Our drive inheritance rate was moderate, and multigenerational cage studies showed quick drive spread to 96-97% of the population. However, the drive failed to reach the whole population due to the formation of functional resistance alleles, despite use of four gRNAs, a strategy that previously was successful at preventing functional resistance. Sequencing showed that these alleles had a large deletion and must have utilized an alternate start codon. The resistance allele had a modest fitness advantage over the drive in a cage study, which could prevent long-term persistence of the drive, especially if cargo genes had an additional fitness cost. Thus, revised design strategies targeting more essential regions of a target gene may often be necessary to avoid such functional resistance, even when using multiplexed gRNAs.

## Introduction

Synthetic gene drive systems bias inheritance to spread throughout a population^1–3^. These systems show the potential of addressing disease transmission vectors, invasive species, and agriculture pests through two primary approaches: population suppression and population modification (also called replacement). Population suppression usually involves disrupting essential genes to reduce fitness and subsequently eliminate the population. For example, by targeting a conserved *dsx* female fertility exon in *Anopheles* mosquitoes, female drive homozygotes became sterile, leading to a decline in the vector population in a laboratory cage experiment^4^. Population modification refers to changing the wild-type population into a transgenic population^5–9^, which can be combined with a cargo gene or disruption of a native gene designed to inhibit transmission of pathogens such as malaria or dengue.

The advent of CRISPR technology has significantly advanced gene drive research. Homing gene drives constructed using Cas9 can precisely cut the wild-type allele on a homologous chromosome at the same genomic site as the drive. Subsequent homology directed repair will use of the drive element as a template, resulting in the copying of the drive element into the wild-type chromosome. This copying mechanism occurs in the germline, resulting in most offspring inheriting the drive, which leads to its rapid propagation throughout the population. This approach have been demonstrated in many organisms including yeast^10–12^, *Drosophila*^7,13–16^, *Anopheles*^4,6,8,9,17^, *Aedes*^18,19^, *Culex*^20^, *Ceratitis capitata*^21^, and *Mus musculus*^22^.

One major issue that homing drives face is resistance alleles that block gRNA binding and Cas9 cleavage^23–26^. These can be present due to natural genomic variation, but they can also form if the Cas9-induced DNA break is repaired by end-joining, which can mutate the target site, forming an indel. This mutation inhibits further recognition by the Cas9-gRNA complex, thereby preventing drive conversion of wild-type alleles. A resistance allele, especially one that carries a fitness advantage over the drive allele, can eventually outcompete the drive in the population. If the gene drive targets a specific gene, then there can be two main resistance allele types: functional and nonfunctional. Functional resistance alleles represent sequences which preserve the function of the target gene. They are expected to have equal or higher fitness than drive alleles under most realistic circumstances. Nonfunctional resistance alleles are more commonly observed because any frameshift mutation or change in important amino acids is expected to disrupt the target gene’s function. These can be removed from the population if the target gene is essential. By using multiple gRNAs targeting nearby sites, high drive conversion efficiency can be preserved and functional resistance alleles can often be prevented because each site must be repaired in a functional manner for the entire allele to remain functional^27,28^.

Suppression drives are themselves designed to disrupt haplosufficient but essential genes, but to target an essential gene with a modification drive, the drive must provide rescue for the gene. In such gene drives, a copy of the gene that has its amino acids recoded to prevent gRNA cleavage (from the drive) serves as a “rescue”. The resistance alleles that were disrupted at the target will be removed from the population, but the drive will still retain high fitness. One successful rescue drive targeted a haplolethal gene with two gRNAs, thus successfully avoiding functional resistance^7^. Because resistance alleles were haplolethal, any offspring inheriting a resistance allele were nonviable, quickly removing resistance alleles from the population. However, engineering drive alleles and rescue elements at haplolethal sites is challenging, even in model organisms^29^. Additionally, high rates of Cas9 cleavage from maternal deposition could form embryo resistance alleles, which could remove drive alleles from the population at significant rates, impairing haplolethal gene-targeting drives. An alternate approach is to target an essential but haplosufficient gene, where only individuals that are homozygous for nonfunctional resistance alleles are nonviable. Though these are easier to engineer and could avoid removal of drive alleles from embryo resistance, they are also substantially slower at removing resistance alleles and will tend to not reach 100% drive allele frequency if there are any fitness costs, though they still are expected to reach 100% drive carrier frequnecy^28^. Thus far, such drives have been made with one gRNA in *Anopheles stephensi*^17^ and *Drosophila melanogaster*^30,31^. All were somewhat successful but still formed functional resistance alleles.

Here, we constructed a homing gene drive targeting the haplosufficient but essential gene *hairy* with four gRNAs. This was based on a similar insertion site to a previously successful CRISPR toxin-antidote gene drive targeting the same gene^32^. Experiments showed only moderate drive conversion efficiency, but the drive was still able to spread to a high level in cage experiments. However, the drive failed to reach 100% carrier frequency due to formation of functional resistance alleles, despite use of multiple gRNAs. This was partially due to use of an alternate start codon site in the functional resistance alleles. The drive also had a moderate fitness cost, which would limit its ability to persist in the face of functional resistance. Thus, additional care is needed for designing drives that target essential regions of a target gene, even with strategies involving multiplexed gRNAs.

## Methods

### Plasmid construction

The HDREh4 drive plasmid was built with restriction digestion, PCR, and Gibson assembly. After transformation in DH5α competent *Escherichia coli,* plasmids were cleaned up by using the ZymoPure Midiprep kit for microinjection. The plasmid was confirmed by Sanger sequencing. Annotated plasmid and genomic DNA sequences are available at (https://github.com/jchamper/ChamperLab/tree/main/Homing-Haplosufficient-Rescue).

### Generation of transgenic lines

The microinjection mix contained 10nM Tris-HCl, 100 μM EDTA solution at pH 8.5, the donor plasmid,HDREh4 (500 ng/ul) serving is its own gRNA source, and the helper plasmid TTChsp70c9^33^ (500ng/ul) to supply Cas9 into *Drosophila melanogaster w*^1118^ embryos. The resulting injected flies were crossed with *w*^1118^. Transgenic offspring with DsRed eyes were selected to identify drive carriers. Homozygous lines were generated by self-crossing following by Sanger sequencing confirmation. The split-Cas9 transgenic line was generated previously^34^.

### Fly rearing and phenotyping

Flies were maintained in the incubator at 25℃ with a 14/10-hour day/night cycle and 60% humidity. Adults flies were anaesthetized using CO_2_ and screened for fluorescence using the NIGHTSEA system for DsRed and EGFP. The eye-specific 3xP3 promoter was used to drive fluorescent protein expression. DsRed and EGFP were respectively used as markers to indicate the presence of the drive allele HDREh4 and the *nanos*-Cas9 allele.

### Drive efficiency test

HDREh4 lines were crossed with the *nanos*-Cas9 line BHDaaN^34^ to generate drive offspring with both DsRed and EGFP fluorescence. These drive offspring were further crossed to *w*^1118^ flies, and their progeny were screened for fluorescence. The percentage of DsRed flies represented the drive inheritance rate.

### Phenotype data analysis

Data were pooled from different to calculate drive performance. However, this pooling approach does not take possible batch effects into account (progeny from a single vial is considered to be a separate batch, see Supplemental Data Sets), which could bias rate and error estimates. To account for batch effects, we conducted an alternate analysis as in previous studies^13,28,35,36^. We fit a generalized linear mixed-effects model with binomial distribution (maximum likelihood, Adaptive Gauss-Hermite Quadrature, nAGQ = 25). This allows for variance between batches, usually resulting in slightly different estimates and an increased standard error. This analysis was performed using R (3.6.1) and supported by packages lme4 (1.1-21, https://cran.r-project.org/web/packages/lme4/index.html) and emmeans (1.4.2, https://cran.r-project.org/web/packages/emmeans/index.html). The code is available on Github (https://github.com/jchamper/ChamperLab/tree/main/Homing-Haplosufficient-Rescue). These alternate rate estimates and errors were similar to the pooled analysis (Supplemental Data Sets).

### Cage study

Flies were maintained in 30 × 30 × 30cm enclosures (BugDorm; BD43030D). In cages A and B, flies with heterozygous drive and homozygous Cas9 alleles were introduced to Cas9 homozygous cage populations at release frequencies of approximately 50% and 30%, respectively, where they could interbreed or mate with Cas9 flies. In our experimental setup for Cage C, we introduced a population where all individuals were heterozygous, each carrying one drive allele and one resistance allele, to initiate generation zero. Cage D was populated with 100% heterozygous flies, with each fly possessing one wild-type allele alongside one resistance allele. Across all cages, eight food bottles were put in each cage for egg collection. After one single day of oviposition, the adult flies were frozen for later phenotyping. In each generation, flies were allowed to grow in the cage for the next 11-12 days (12 days if over 50% of the pupa had not eclosed), after which new bottles were placed in cages for 24 hours to collect eggs as next generation.

### Deep sequencing for resistance alleles

Cages A and B were maintained after the end of the experiment until the 16th generation. We collected all fruit fly samples from the 16th generation of cages A and B, and all fruit fly samples from the 2nd, 6th and 10th generations of cages D for deep sequencing. Flies genomic DNA were extracted with DNAzol (Thermo Fisher Scientific). Primers P5RD1 and P7RD2 (see Supplemental Information) were used to amplify a 514 bp fragment covering gRNA target sites, and the PCR products were cleaned up for deep sequencing by Novogene (using a NovaSeq 6000, PE250). All reads are available on GitHub (https://github.com/jchamper/ChamperLab/tree/main/Homing-Haplosufficient-Rescue). Other DNA sequences were obtained via Sanger sequencing and analyzed using the ApE software^37^.

### Fitness cost inference framework

To assess drive fitness costs, we used our maximum likelihood inference framework. Similar to previous studies^7,13,33,38^, we analyzed our homing rescue drive. The maximum likelihood inference method is implemented in R (v. 4.0.3) and is available on GitHub (https://github.com/jchamper/ChamperLab/tree/main/Homing-Haplosufficient-Rescue). In this model, we assume a single gRNA at the drive allele site. Each female randomly selects a mate, though the relative chance of drive males being selected is equal to their fitness for mating success models. The number of offspring generated is proportional to the fitness of drive females for fecundity fitness costs. In the germline, wild-type alleles in drive/wild-type heterozygotes can be converted to either drive or resistance alleles, which can be functional or nonfunctional. The genotypes of offspring can be altered if they have a drive-carrying mother and if any wild-type alleles are present. These alleles can be converted to resistance alleles at the embryo stage. Viability fitness costs can then eliminate offspring (with survival equal to their relative fitness), and offspring with two nonfunctional resistance alleles are always nonviable.

## Results

### Construction of the haplosufficient rescue drive

In this study, we built a homing drive construct targeting the *hairy* gene with rescue in *Drosophila melanogaster*. The drive copies itself into wild-type alleles in the germline, biasing inheritance (Figure 1). Resistance alleles can also form in the germline or later in the embryo due to maternal deposition of Cas9 and gRNA. The target gene is haplosufficient but essential. Thus, individuals with two nonfunctional resistance alleles will be nonviable. Even one copy of wild-type, drive, or functional resistance alleles is sufficient to ensure viability.

**Figure 1.**
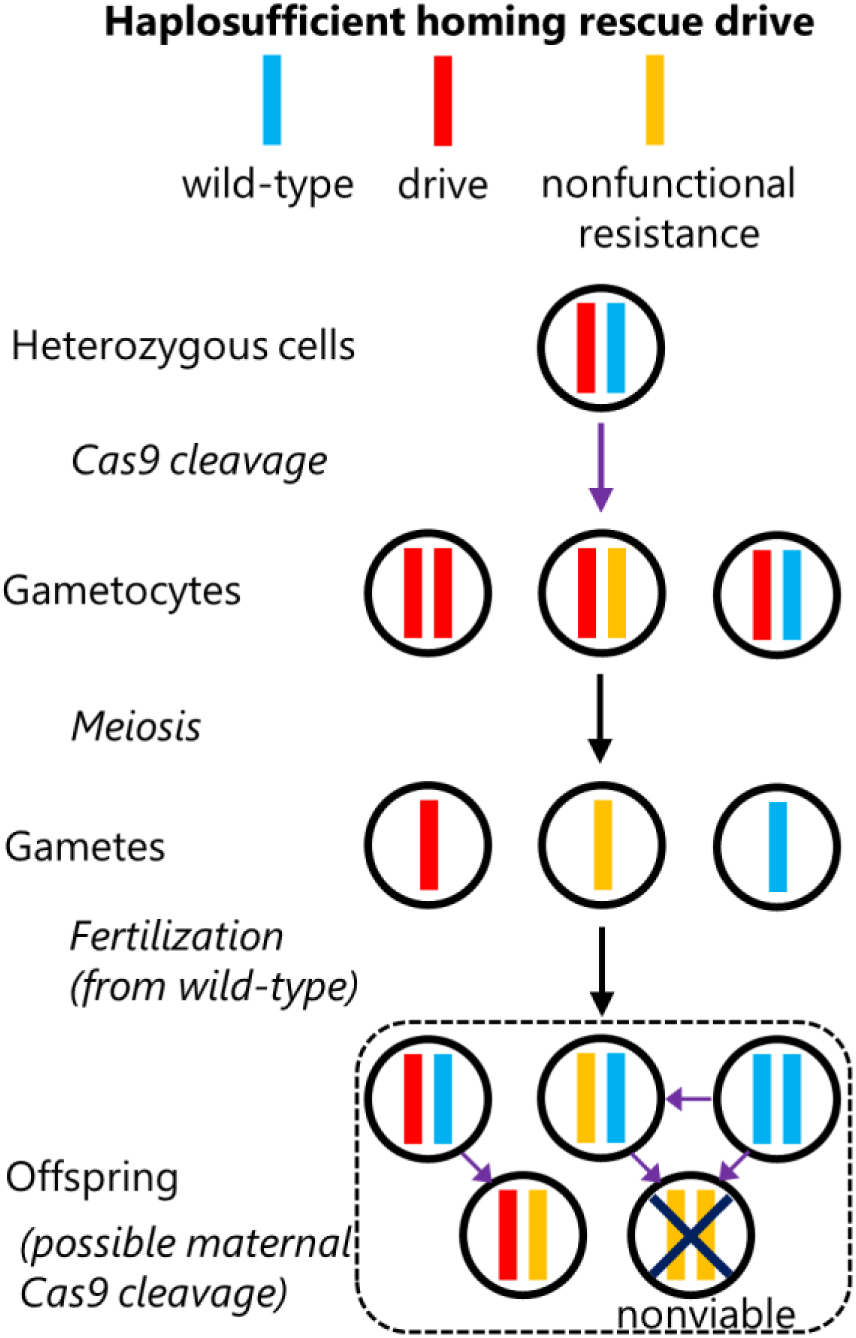
SIT. The homing drive can cut the wild-type allele in the germline, leading to successful drive conversion or resistance allele formation. After meiosis and fertilization, additional wild-type alleles can be converted to resistance alleles due to maternal deposition of Cas9 and gRNA. Because the drive targets the haplosufficient but essential hairy gene, homozygotes for nonfunctional resistance alleles are nonviable. Functional resistance alleles can form whenever nonfunctional resistance alleles can in this chart, but one functional resistance allele is sufficient to ensure viability.

Specifically, we modified a previously constructed CRISPR toxin-antidote drive^32^, changing the gRNAs and slightly adjusting the recoded portion and left homology arm (Figure 2). The construct thus carried a recoded version of *hairy*, which could not be cleaved by Cas9/gRNA and thus served as a rescue copy of the *hairy* gene, allowing drive carriers to remain viable. The construct also expressed the fluorescent protein DsRed driven by the 3xP3 promoter, which could be used for eye-specific phenotyping. The U6:3 promoter was used to drive four different gRNAs which were linked by tRNAs that would be spliced out, allowing expression of all gRNAs from the same promoter. These multiplexed gRNAs targeted different sequences in the coding sequence of *h* gene. They were closely spaced to maintain drive conversion efficiency while reducing the possibility of producing functional resistance alleles. This drive was designed as a split drive for biosafety and was tested with a *nanos*-Cas9 allele located on a separate chromosome that was constructed previously^34^.

**Figure 2.**
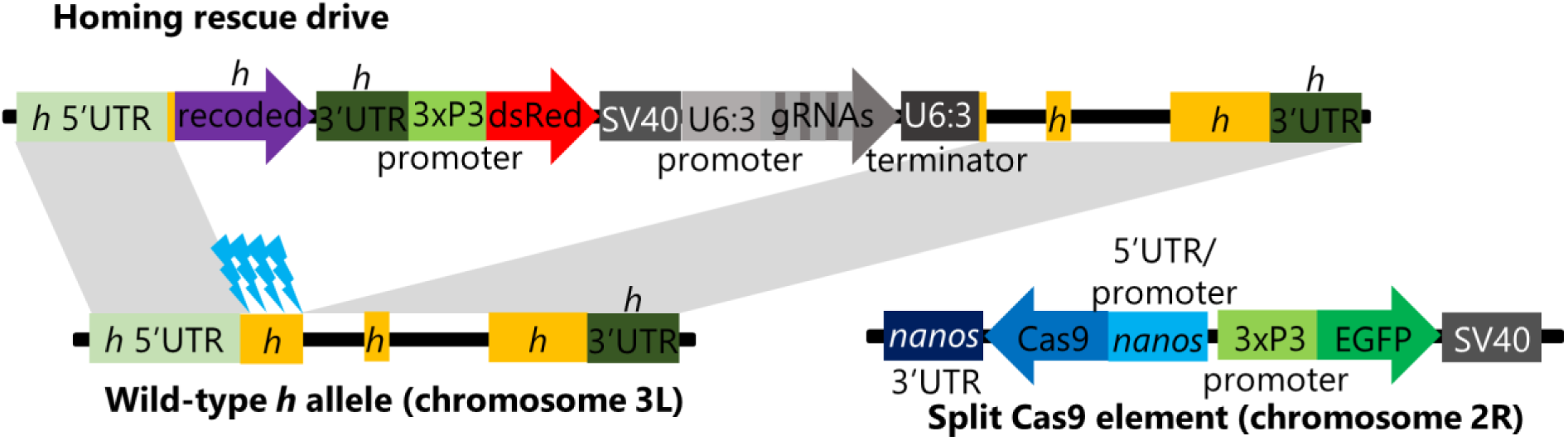
Drive element schematic. The gene drive was inserted into the first exon of *hairy*. It contains a recoded version of the *hairy* condign sequence followed by the native *hairy* 3’UTR, which together serve as a rescue version of *hairy* that cannot be cleaved by the drive’s gRNAs. Next is a DsRed, using the 3xP3 promoter and SV40 terminator for strong expression in the eyes, allowing easily phenotyping of drive individuals. The gRNA gene using the U6:3 promoter and includes tRNAs between the gRNAs, thus allowing separation of the gRNAs from a single transcript. The gRNA all target the first exon of *hairy*. On a separate chromosome, Cas9 driven by the *nanos* promoter provides for Cas9 expression in the germline, enabling drive conversion and low somatic expression, but also high maternal deposition of Cas9. *h* – *hairy* gene.

### Super-Mendelian inheritance of the homing rescue drive

To assess drive conversion efficiency, we crossed homozygous or heterozygous drive females with Cas9 homozygous males. Their drive offspring, which expressed both red and green fluorescence, were then selected to cross with *w*^1118^ flies. Offspring from these crosses were collected and phenotyped (Figure 3, Data Set S1). The results showed a 68.6% drive inheritance rate in progeny of gene drive males and 67.9% drive inheritance in gene drive females. The drive inheritance rates of male and female drive crosses were both significantly higher than 50% expected under Mendelian inheritance (*P*<0.0001, binomial exact test).

**Figure 3.**
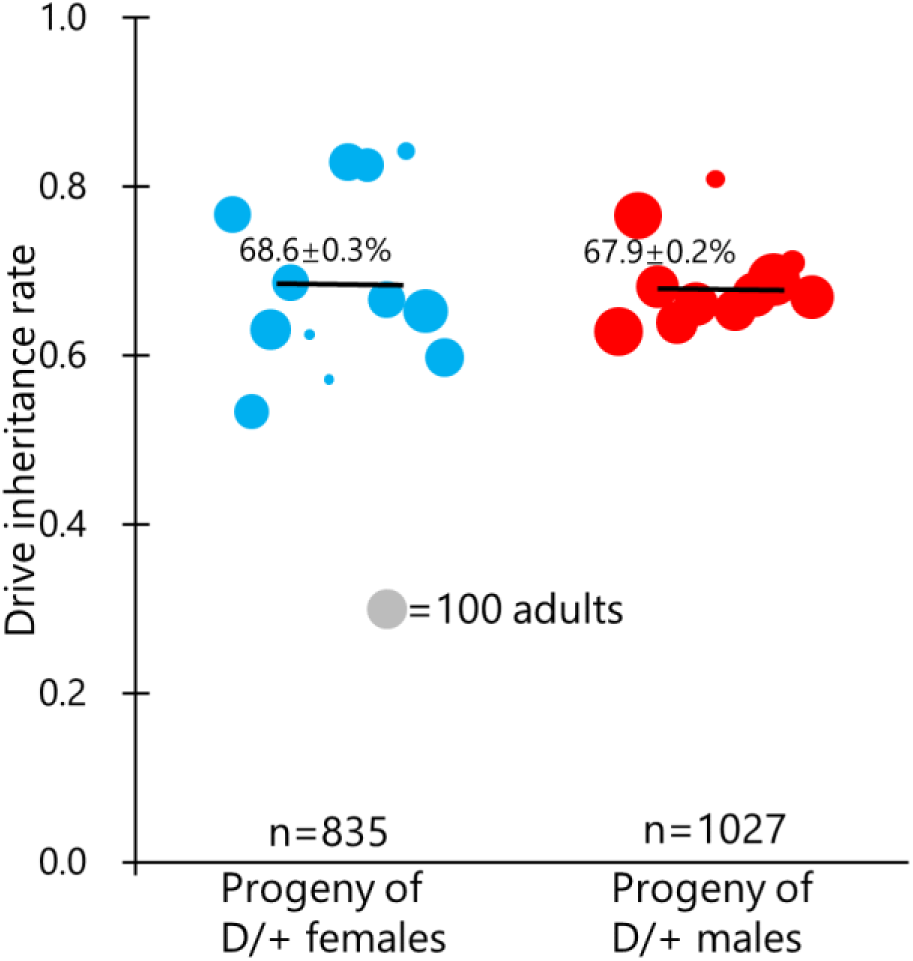
Drive performance. The drive inheritance rate among progeny of drive heterozygotes with one copy of Cas9 is shown. Each dot represents progeny from a single drive individual in one vial, and the size of each spot shows the number of individuals phenotyped. The mean inheritance rate (± standard error of the mean) is shown.

### Multigenerational cage experiments

To test whether homing rescue drives can spread through and modify a population, we set up cages by releasing drive carriers into cage populations at a frequency of either 50% (cage A) or 30% (cage B). The results showed that our drive was able to spread and modify the population in both cages (Figure 4A, Data Set S2). In cage A, the drive carrier frequency reached a peak at 97%, but then began a slow decline. In cage B, the drive carrier frequency increased to approximately 96%. The cage populations were maintained but not phenotyped past generation 10 (see below).

**Figure 4.**
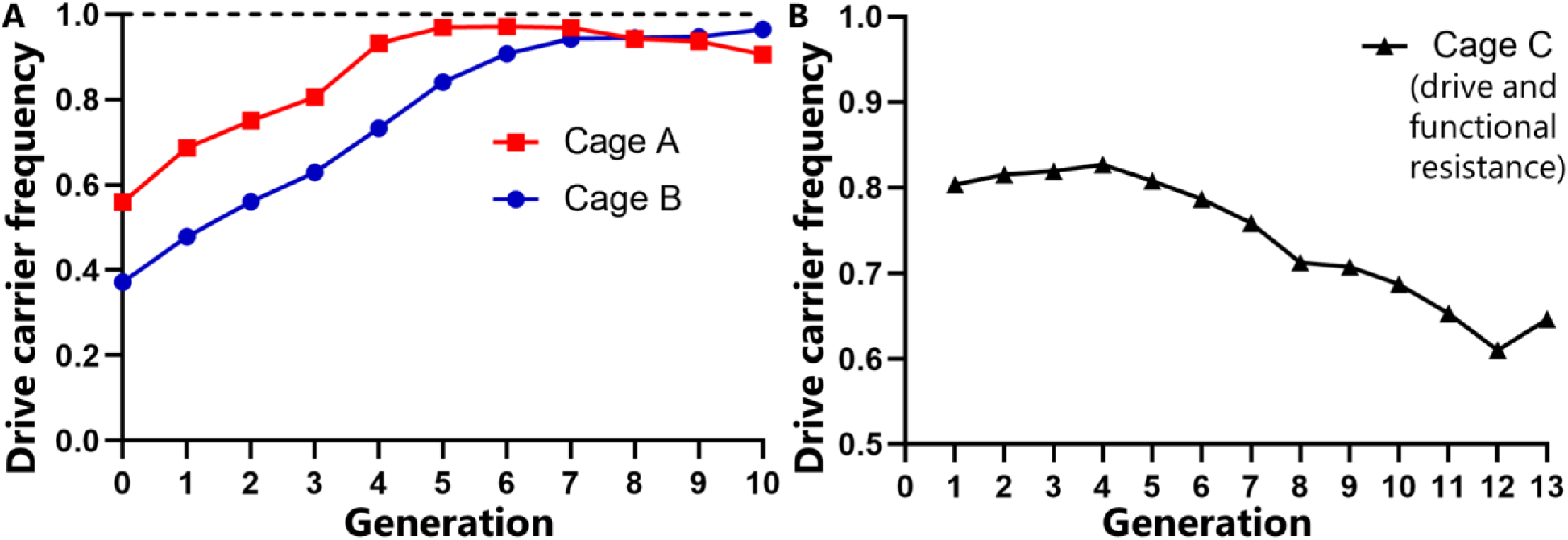
Cage studies. (**A**) Females homozygous for Cas9 were allowed to mate with either Cas9 homozygous males or drive heterozygous males (that were also homozygous for Cas9). The progeny were then considered to be “generation 0” for the cage. All individuals were phenotyped for ten discrete generations to track drive carriers. (B) Non-drive flies from generation 13 of cages A and B were crossed together for two generations and then crossed to drive homozygotes to form generation 0, all of which were drive/resistance allele heterozygotes. The drive carrier frequency was then followed for several discrete generations. See Figure S1 for population sizes.

### Analysis of functional resistance alleles in cage experiments

We reasoned that even a weak drive with low cut rates should be able to cut most wild-type alleles by generation 10, so only functional resistance alleles could adequately explain why the drive failed to reach 100% carrier frequency. To confirm this, we collected flies without the drive from both cages at generation 10 and sequenced the target site, while performing additional sequencing to detect undesired homology-directed repair in which only the rescue element would be copied. The results from eight flies showed no instances of undesired homology-directed repair, but all had indels at gRNA target sites, representing resistance alleles. Interestingly, gRNA 1 targeted the translation start site of *hairy*, eliminating the start codon and any Kozak sequence that was present.

However, a potential alternative translation start site was found downstream in the first exon and in-frame of the normal amino acid sequence. This likely maintained the protein function of *hairy* in these non-drive mutant flies.

Deep sequencing of the drive target site of all individuals from generation 16 of these cages revealed a similar pattern (Figure 5, Data Set S4). Nearly all of the most common resistance alleles were in-frame for either the normal start codon or the second methionine codon in cases where the first was deleted. Their high frequency suggests that they had a fitness advantage over other resistance alleles and were thus functional. The low sequence diversity at individual cut sites suggests possible use of microhomology-mediated repair pathways. Despite 16 generations of potential Cas9 cleavage, many wild-type sequences remained, and even for the functional resistance alleles, many still had uncut sites (especially at the second gRNA) that could potentially allow future drive conversion. This low cut rate is consistent with the low ∼35% drive conversion efficiency.

**Figure 5.**
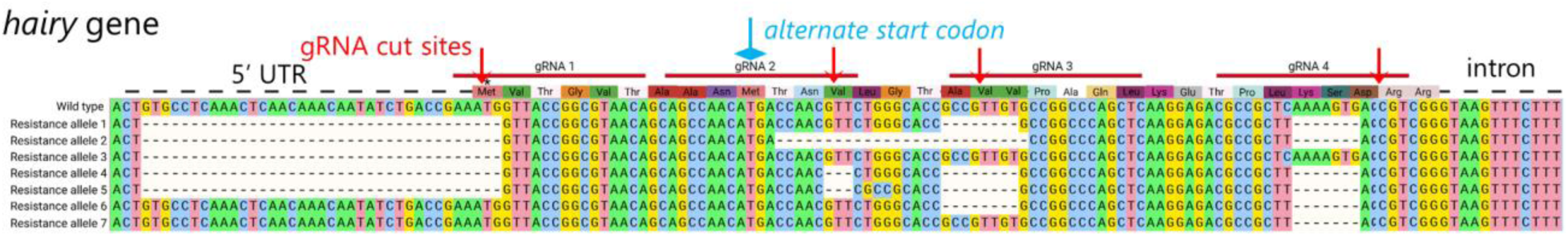
Functional resistance alleles. Deep sequencing was conducted for the drive cage experiments, and the seven most common alleles were identified. Each of these is in frame, either for the original start codon or an alternate start codon. Some of these resistance alleles are complete, while others retain one or more gRNA target sites that may eventually be cut, rendering them nonfunctional or converting the entire allele to a drive allele.

Further analysis suggested why in-frame mutations may consistently produce functional resistance alleles in exon 1 of the protein. *hairy* lacks secondary structure in this region^39^, which is flexible, thus reducing the impact of mutations on the proteins overall structure. Furthermore, the protein appears to lack a signal peptide^40^, which can increase the importance of the first few amino acids of other proteins. Finally, because *hairy* is haplosufficient, changes in the expression level from deletions of the Kozak sequence and alternate start codon may not have a drastic effect on an organism’s health.

To evaluate the fitness of the resistance alleles identified in cages A and B, we established two additional cages, C and D. Cage C was designed to assess the relative fitness between the resistance allele and the gene drive allele. We initiated the experiment with a population of 100% heterozygous flies (generation zero), each carrying one gene drive allele and one allele derived from non-drive flies from generation 13 of cage A and B (most of which should have been functional resistance alleles or wild-type alleles). The drive showed a progressive decline in frequency (Figure 4B), suggesting that functional resistance alleles had a modest fitness advantage over the drive.

Because the resistance allele we detected had major deletions in potentially important protein functions, we were curious to see whether it might have a fitness disadvantage compared to wild-type alleles. To investigate this, we set up Cage D, which was set up similarly to Cage C, but crossing the non-drive Cage A/B flies to wild-type flies instead of drive homozygous flies.

Because there was no visible phenotype in these cages, we collected and sequenced all individuals using deep sequencing from generation 2, 6, and 10 (Data Set S4). The wild-type allele frequency at these generations was 67%, 65%, and 77%, respectively, suggesting a potential small fitness advantage of the wild-type allele over the drive allele.

### Maximum likelihood analysis of cage experiments

To evaluate the fitness of the drive allele in the experimental cage studies, we applied an established maximum-likelihood approach tailored for homing gene drives targeting haplosufficient genes. We initially analyzed Cage C, comparing the drive and resistance alleles. Using the simplifying assumption of a starting population of only drive and resistance alleles, we found that the best fit model had a combined female fecundity and male mating success fitness of 0.9 (where 1 is equal to wild-type) (Table S1A), representing the fitness of drive homozygotes, with multiplicative fitness per drive allele.

The 95% confidence interval ranged from 0.81 to 0.99, which is broadly consistent with our previous studies of modification drives^7,35^. The moderate effective population size indicated that the model was a decent fit.

Our combined analysis of Cages A and B assumed a single guide RNA and a male gene drive conversion rate of 35.4%, based on our data for individual crosses. The female drive conversion rate was dynamically adjusted to produce a drive inheritance rate of 68.6%, even with varying germline and embryo resistance rate. We used our fitness estimate of 0.9 from cage C as a fixed parameter and varied the functional and nonfunctional resistance allele formation rates, allowing functional resistance alleles to represent no more than 1/3 of total resistance alleles because they must still be in-frame. Because resistance allele formation rates are low based on our low drive conversion and sequencing, and embryo cut rates usually tend to be substantially lower than germline cut rates based on our previous studies with *nanos*-Cas9^7,13,23,27,35^, we fixed the embryo resistance level at half of the germline resistance allele formation rate. Male and females were assumed to have identical germline resistance allele formation rates. Under these conditions, the best fit model (Table S1B) had a total germline resistance allele formation rate of 64%, with the functional resistance allele formation rate equal to 9% of the nonfunctional resistance rate.

However, a model with functional resistance fixed at 1/3 of the total was nearly as good and perhaps closer to reality, considering the relatively low overall cut rates of this drive and the low drive conversion efficiency. These model fits were somewhat poorer than Cage C based on the effective population size estimate, perhaps due to the more complex situation.

## Discussion

In this study, we investigated the possibility of developing a homing rescue gene drive targeting *hairy*. Although we used four gRNAs, our results showed that this did not significantly reduce the rate of functional resistance allele formation. However, the drive still reached a high carrier frequency in the population.

In a previous models^28^, four gRNAs was more than sufficient to avoid functional resistance alleles (indeed, even two gRNAs was sufficient in laboratory population experiments^7,27^). However, this relies on the assumptions that most in-frame mutations will still disrupt protein function and that a frameshift between two gRNAs would be sufficient to disrupt the protein (by mutating several amino acids, even if a later indel restores the reading frame). We unexpectedly found that *Drosophila melanogaster* can survive even after the loss of a large fragment in the first exon of *hairy*, with mutations at all four gRNA sites, as long as there is no frameshift in the latter part of the protein. This is likely possible because the amino acid sequence from the first exon is nonessential, at least in laboratory conditions. The protein itself is predicted to have no secondary structure in this portion, supporting the notion that it is a less essential part of the protein. We also several different functional resistance allele sequences that must have formed at high rates, further supporting the notion that at least many mutations will produce functional resistance, even if regions between the gRNA cut sites are out-of-frame.

Interestingly, we observed a novel form of resistance allele in this study. Though we targeted and eliminated the start codon of *hairy*, which we believed would itself usually ensure that any resistance alleles were nonfunctional, we found many resistance alleles remained functional despite this deletion. We suspect that this may be because of a second start codon in the first exon that avoided disruption. Though the original Kozak sequence would be less able to support this second start codon, the haplosufficient nature of this gene may mean that an allele can be an effective functional resistance allele even if has potentially reduced expression compared to the wild-type allele. Our deep sequencing suggests that it might have a small disadvantage compared to wild-type, but more importantly, our maximum likelihood analysis suggests that it still has a fitness advantage over the drive allele. For a complete drive, the fitness advantage may be slightly greater^35^, and a cargo gene may further reduce drive fitness, limiting the long-term persistence of the drive allele, even though it reached a high frequency in the population.

This target site was chosen based on the insertion site of a previously constructed CRISPR toxin-antidote drive that had high efficiency^32^. This was because it targeted the middle of the *hairy* coding sequencing, avoiding functional resistance with only two gRNAs in this more essential region of the protein. While such a verified insertion site (with gRNAs that are known to be active for transformation) may be convenient, our study shows that even when targeting coding sequence, care should be taken to find essential regions of the target gene, which can often itself be sufficient to prevent functional resistance in laboratory populations^4^. Even targeting a highly conserved start codon may be insufficient if there are alternate candidates.

At ∼35%, the drive conversion efficiency of our drive in this study was considerably lower than most previous drives in *D. melanogaster* that utilized the *nanos* promoter for Cas9, despite some of these gRNAs being useful for transforming the CRISPR toxin-antidote drive^32^. The reasons for this are somewhat unclear considering that the same gRNA promoter and multiplexing system were used. While the gRNAs themselves may have been less active on average, the genomic location may have also played a role. However, our cage studies show that for modification drive (unlike suppression drive^13^), modest drive efficiency won’t necessarily have a negative impact on the ultimate outcome of the drive, merely somewhat slow it down.

In summary, despite modest drive conversion efficiency, our rescue drive targeting the haplosufficient gene hairy reached high frequency in cage studies. Yet, the drive was thwarted by formation of functional resistance alleles, despite targeting of the start codon and use of four gRNAs. In the long-term, the drive’s modest fitness could would allow these resistance alleles to outcompete the drive. In order to minimize the development of functional resistance alleles, future drives should target regions of a gene that are more important for the protein’s structure or function, thus preventing functional resistance in the context of a gRNA multiplexing strategy.

## Acknowledgements

This study was supported by laboratory startup funds from the Center for Life Sciences and the NSFC Overseas Youth Fund.

## Supplemental Information

### Deep sequencing primers

**Figure S1.**
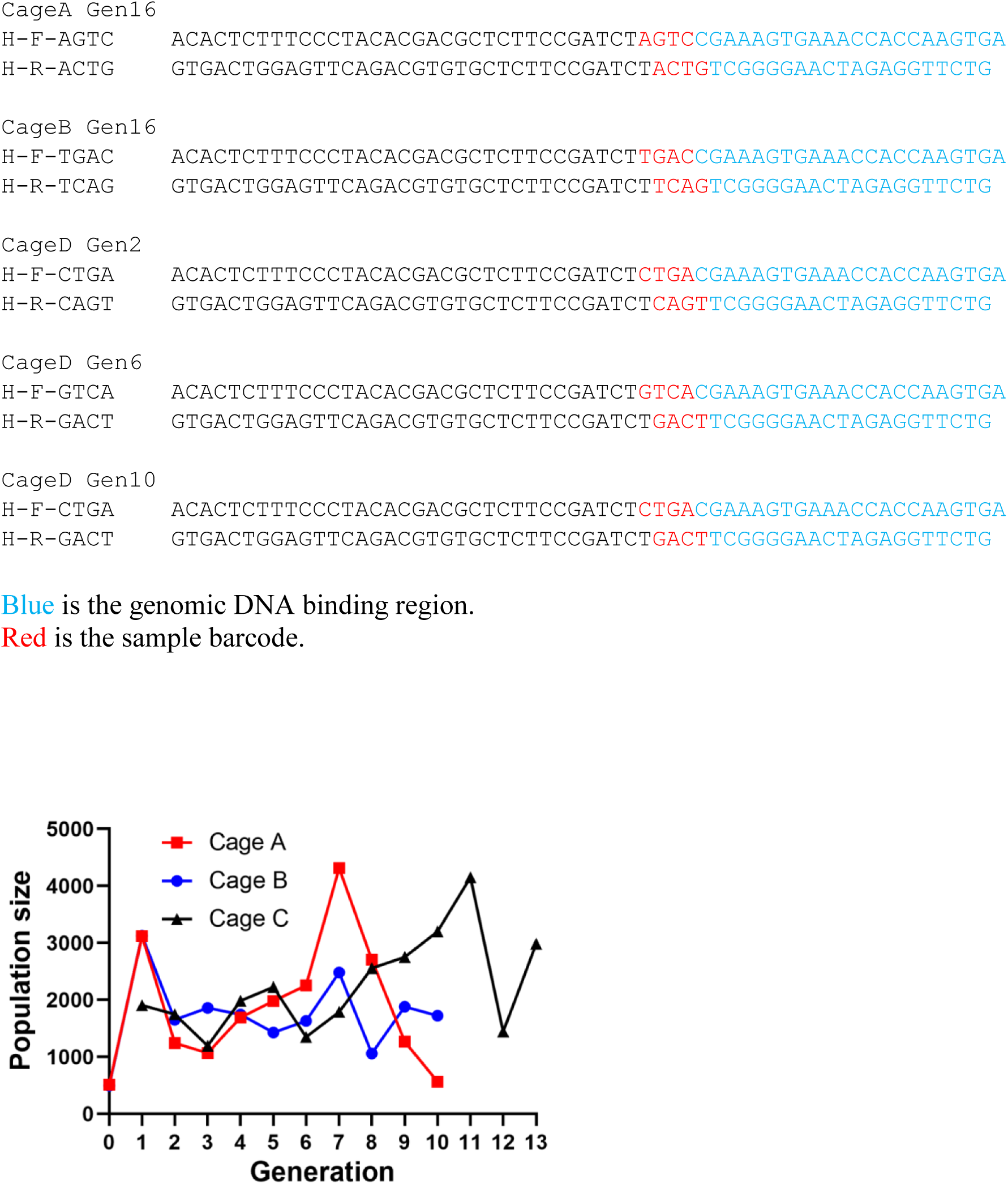
Cage population sizes. The chart shows the number of individuals for each generation of the cages from Figure 4.

**Table S1.**
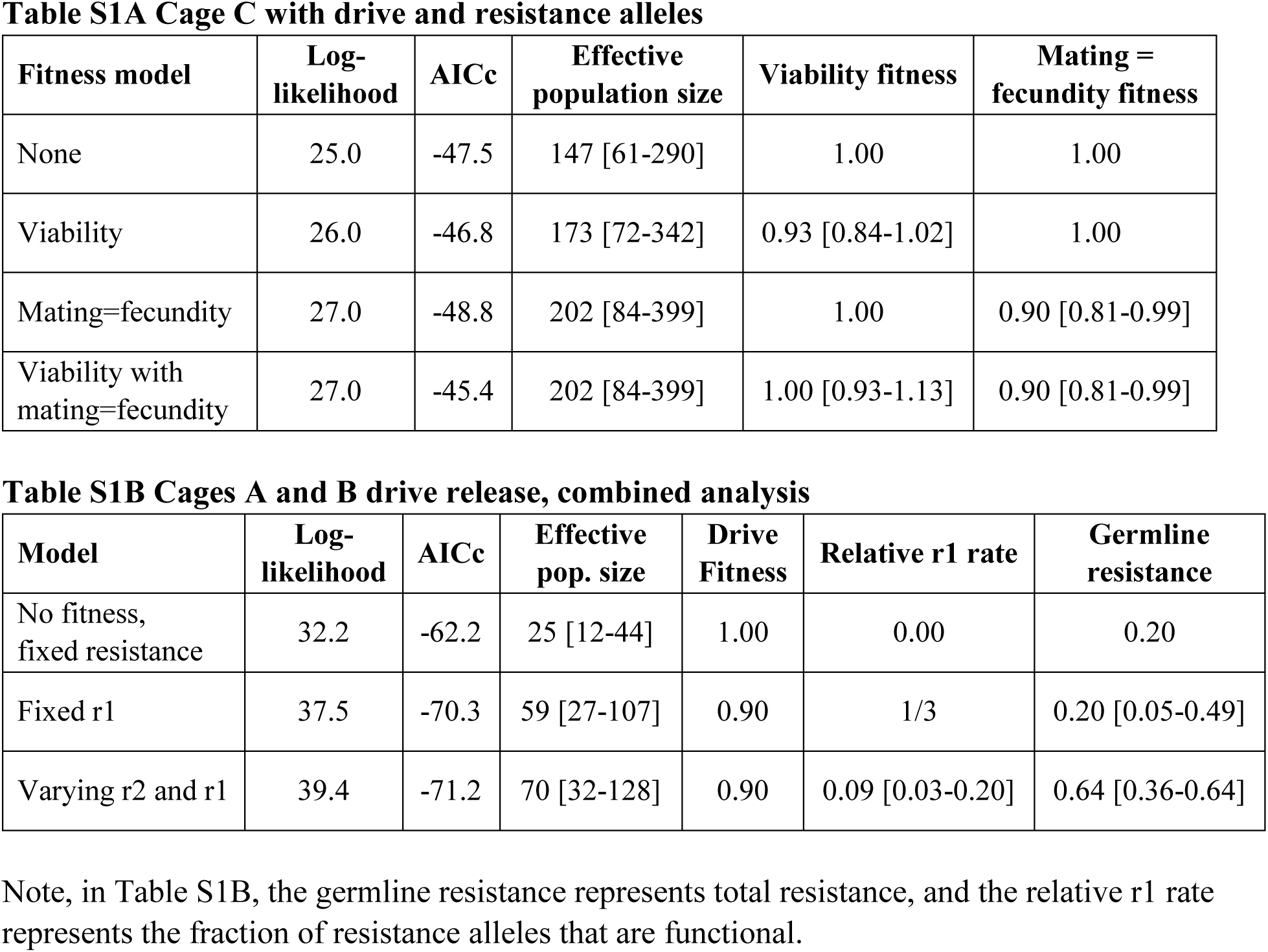
Maximum likelihood parameter estimates from cage populations. Fitness values are for drive homozygotes (multiplicative fitness per allele). 1 is equivalent to wild-type. [Brackets] show 95% confidence intervals. All models have an effective population size parameter. Log-likelihood: shows relative probability (higher values indicate a better model fit) AICc: Akaike information criterion, corrected (low values indicate a better match of the model without overfitting).

